# Sustained polyphasic sleep restriction abolishes human growth hormone release

**DOI:** 10.1101/2023.06.20.542775

**Authors:** Yevgenia Rosenblum, Frederik D. Weber, Michael Rak, Zsófia Zavecz, Nicolas Kunath, Barbara Breitenstein, Björn Rasch, Marcel Zeising, Manfred Uhr, Axel Steiger, Martin Dresler

## Abstract

Voluntary sleep restriction is a common phenomenon in industrialized societies aiming to increase time spent awake and thus productivity. We explored how restricting sleep to a radically polyphasic schedule affects neural, cognitive, and endocrine characteristics. Ten young healthy participants were restricted to one 30-min nap opportunity at the end of every 4 hours (i.e., 6 sleep episodes per 24 hours) without any extended core sleep window, which resulted in a cumulative sleep amount of just 2 hours per day (i.e., ∼20 min per bout). All but one participant terminated this schedule during the first three weeks. The remaining participant (a 25-year-old male) succeeded to adhere to a polyphasic schedule for 5 weeks with no apparent impairments in cognitive and psychiatric measures except for psychomotor vigilance. While in-blood cortisol or melatonin release pattern and amounts were unaltered by the polyphasic as compared to monophasic sleep, growth hormone seemed almost entirely abolished (>95% decrease), with the residual release showing a considerably changed polyphasic secretional pattern. While coarse sleep structure appeared intact during polyphasic sleep, REM sleep showed decreased oscillatory and increased aperiodic EEG activity compared to monophasic sleep. Considering the decreased vigilance, abolished growth hormone release, and neurophysiological changes observed, it is doubtful that radically polyphasic sleep schedules can subserve the different functions of sleep to a sufficient degree.

## Introduction

From an evolutionary point of view, spending several hours each day in a non-responsive, vulnerable state seems strikingly non-adaptive. Sleep thus serves several vital functions to overcompensate for this disadvantage. In modern societies, however, voluntary sleep restriction is a common phenomenon, aimed to increase time spent awake while preserving daytime productivity.

Over the last decades, different polyphasic sleep schedules have been advocated in an attempt to enhance the efficiency of sleep and thus potentially reduce cumulative time needed to spend asleep. A polyphasic sleep schedule that achieved some notoriety in internet communities is the *“Uberman”* schedule, characterized by six 20-min sleep episodes spaced evenly across the 24-h day without an extended core sleep period, resulting in a cumulative sleep amount of only 2 hours per day.

There is an appeal of simple behavioral strategies to reduce sleep need, as well as anecdotes that claim polyphasic sleep schedules could improve productivity, memory, mood, dream recall, lucid dreaming frequency, and even longevity. Considering this appeal and potential benefits, empirical evidence on this with polyphasic sleep cycles beyond biphasic siesta/nap habits is surprisingly sparse (Stampi et al., 1989; 1992; Weaver et al., 2021). However, polyphasic sleep schedules might have a detrimental effect on cognitive functioning, and physical and mental health, in line with abundant research on sleep deficiency, which is inherent in most polyphasic schedules (McCoy et al., 2011; Weaver et al., 2021). Given that sleep loss adversely affects almost every process and organ in the body, there arises a question of whether a polyphasic sleep schedule is also harmful to one’s health. To address this question, here, we explore the impact of a sustained polyphasic sleep schedule on cognitive, endocrine, and neural activity of healthy volunteers who decided to change their sleep habits to the *Uberman* schedule. We hypothesized that this schedule will adversely affect all tested domains.

## Methods

### Participants

Twenty healthy university students participated in the study. Ten healthy young adults (mean age = 23.9 ± 2.4, range: 21–28, one female) volunteered to participate in the polyphasic sleep condition. The participants contacted the sleep laboratory of the Max Planck Institute of Psychiatry, Munich, Germany, as all of them decided to voluntarily change their sleep habits to a radically polyphasic schedule. Eight out of 10 participants agreed to undergo a 24-h in-depth monitoring that included polysomnography, blood sampling and cognitive testing, whereas 2 participants restricted their participation to questionnaires. Importantly, all but one participant terminated the polyphasic sleep schedule prematurely within the first three weeks without a second in-depth laboratory appointment (see Fig. S1A). One participant (a 25-year-old male) terminated the polyphasic sleep schedule after five weeks, however, agreed to a second in-depth laboratory appointment before termination. Ten age-matched university students were recruited and served as a control group (mean age = 23.8 ± 1.9, range = 20–27, all males). They stayed on a monophasic sleep schedule (∼8 h of sleep per day) throughout the study period. The study was approved by the Ethics committee of the University of Munich. All participants gave written informed consent. Because polysomnography was not recorded on the original control sample, we performed additional analyses of sleep architecture and spectral power of brain signals on another large sample of participants described elsewhere (Ackermann et al., 2015). From this sample, we chose only age- and gender-matched participants (n = 97, mean age = 24.7 ± 1.4, range = 23–27, all males) and used it as a sufficiently powered reference dataset of normal sleep using the same recording technology.

### Study design

The originally planned length of the study was 12 weeks, out of which, the polyphasic participants were asked to stay 4 weeks on a monophasic sleep schedule as a baseline and then switch to a polyphasic sleep schedule for 8 weeks. Throughout this period, participants filled out daily sleep logs which were validated by actigraphy. Weekly, they filled out questionnaires about their mood and subjective cognitive performance. At the end of the baseline period and planned again at the end of the polyphasic period, participants underwent in-laboratory testing sessions with 24-h physiological monitoring, including polysomnography, blood sampling, and cognitive assessments.

Only one polyphasic participant completed the full study: he underwent all the assessments twice, before and towards the end of the polyphasic sleep schedule. The 10 control participants fulfilled all the assessments as planned.

Sleep architecture, spectral EEG power, and hormone characteristics of the polyphasic participant were compared using a within-subject design, from the monophasic baseline session to 5 weeks later, the polyphasic session (Fig. 1–3). In addition, the sleep architecture of this participant was compared to that measured using the reference dataset (n = 97). The cognitive and affective characteristics of the polyphasic participant were compared to those of the monophasic control participants (n = 10, Fig. 4).

**Figure 1.**
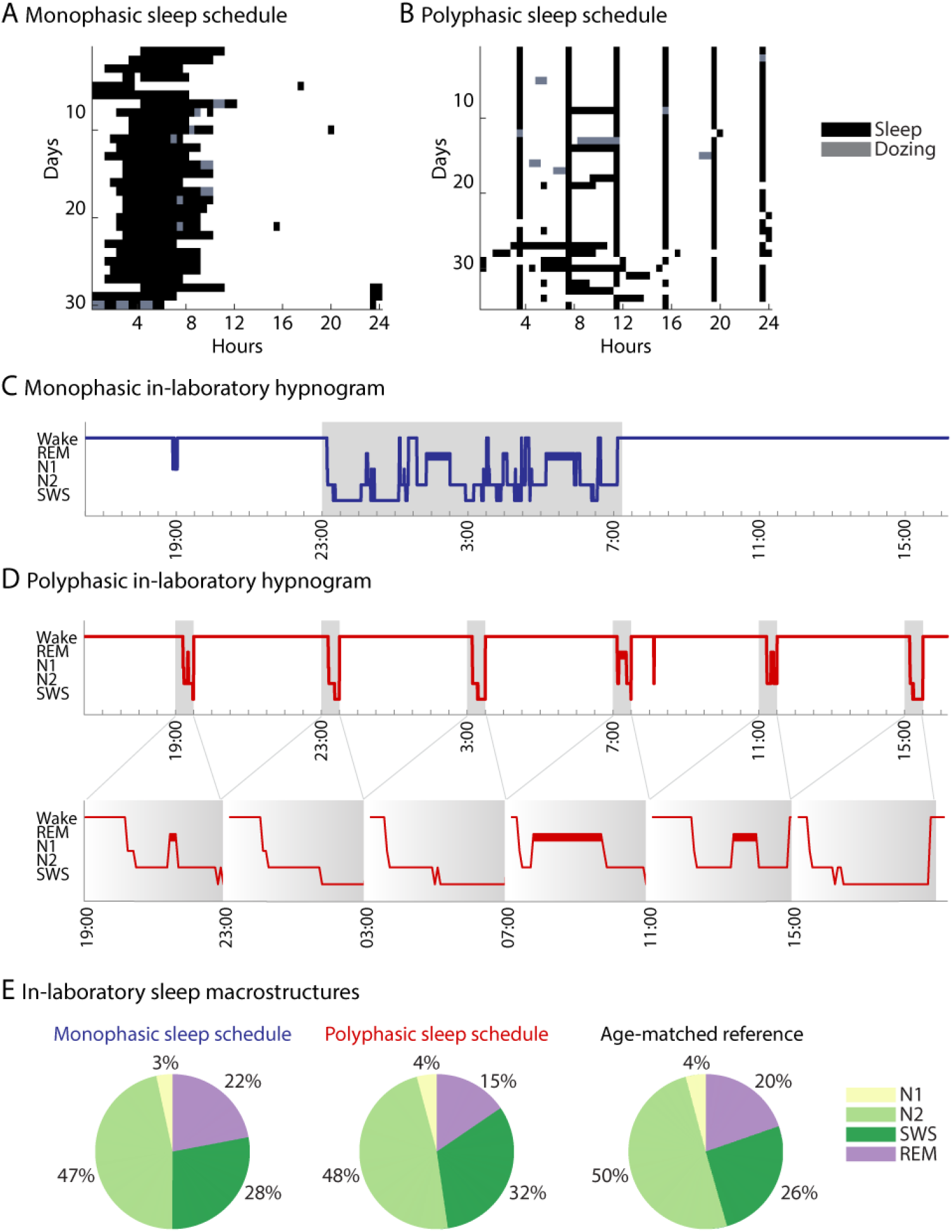
Monophasic and polyphasic sleep schedules. Sleep diaries before (A) and after (B) switching to the polyphasic *“Uberman”* sleep schedule, when the participant slept 20-min every 4 hours (six episodes per day). Black shading indicates sleep, whereas gray indicates dozing. During the polyphasic period (B), the participant succeeded in strictly adhering to the *Uberman* schedule for 22/35 days of the experiment with a median cumulative sleep time of 3 h/day. For 13/35 days, he showed a schedule with an extended core sleep period of ≥4 hours. Sleep hypnograms of the polysomnography monitored in-laboratory monophasic (C) and polyphasic (D) sleep schedules reveal comparable sleep architecture. Gray shading indicates sleep periods that were used to calculate the macrostructure of the specific sleep sessions. (E) The polyphasic sleep schedule (middle) shows a 7% decrease in the proportion of REM sleep in favor of SWS compared to the monophasic schedule (left). The polyphasic and monophasic sleep architecture of the participant is comparable to the sleep architecture measured in the reference dataset (right, n = 97). N2: non-rapid eye movement stage 2, SWS: slow-wave sleep, REM: rapid eye movement.

### Polyphasic sleep

The polyphasic participants adhered to the *“Uberman”* sleep schedule with one 30-min nap opportunity at the end of every 4 hours without any extended core sleep period, resulting in a planned cumulative sleep amount of only 2 hours per day with 6 sleep episodes of about 20 min of sleep. Both monophasic and polyphasic sleep schedules were monitored for 4 weeks before the switch to the polyphasic schedule and during the polyphasic sleep period. Participants filled out sleep logs to indicate when they slept, and in addition, wore an actigraphic device on their ankles (Actiwatch, Cambridge Neurotechnology, Cambridge, UK). At the end of the experiment, all polyphasic participants were interviewed about the impact of this sleep schedule on their life and about potential reasons for terminating the polyphasic sleep schedule (Fig. S1 B).

### Polysomnography

The participant spent two 24-h periods in the sleep laboratory, before the switch to the polyphasic sleep schedule and at the end of the experiment. Polysomnography was recorded for a full day; it included 3 EEG channels (F4, C4, O2 channels referenced to left mastoid), electrooculogram, and mental/submental electromyogram (SOMNOwatch; Somnomedics, Randersacker, Germany). The device was applied in the laboratory between 16:00 and 18:00. Sleep was scheduled from 23:00 (lights off) to 07:00 (lights on) during the monophasic condition and every 4 hours during the polyphasic condition (*“Uberman”* sleep schedule). During the day, most of the time the participant stayed in the laboratory.

We compared neural activity of the polyphasic and monophasic sleep intra-individually within a single participant, and inter-individually with age-matched controls from the reference dataset (n = 97; for details see Ackermann et al., 2015), which was recorded using the same polysomnography system (SOMNOwatch; Somnomedics, Randersacker, Germany).

Sleep data was scored according to standard criteria (AASM, Iber, 2007). To further investigate the sleep architecture during different sleep schedules, we used an automatic sleep-scoring algorithm to gain probabilities of scoring certainty for each sleep stage (Abou Jaoude et al., 2020). It is important to note that we did not use the scoring of the algorithm, only the probabilities of its predictions (Fig. S3).

EEG data was re-referenced to the average of all electrodes and downsampled to 250 Hz. The epochs with artifacts were detected by a visual inspection and rejected from further analysis. Total spectral power for each 30-s epoch for each channel was calculated and then differentiated into its oscillatory and aperiodic (i.e., fractal, 1/f, scale-free) components using the Irregularly Resampled Auto-Spectral Analysis (Wen & Liu 2016). To implement the algorithm, we used the Fieldtrip toolbox (Oostenveld et al. 2011) as described elsewhere (Rosenblum et al., 2022; 2023 a). Briefly, total power and its aperiodic component were calculated using the *ft_freqanalysis* function with *cfg.method = ‘mtmfft’* and *cfg.method = ’irasa*’, respectively. The aperiodic power component was then transformed to log-log coordinates by standard least-squares regression. To estimate the power-law exponent β (the rate of the spectral decay), we calculated the slope of the aperiodic component, a neurophysiological marker of excitation-to-inhibition ratio and arousal states (Gao et al., 2017, Lendner et al., 2020). We calculated the slopes of the aperiodic component in the low (1–30 Hz) and high (30–48 Hz) frequency bands for each sleep stage separately and averaged them over all electrodes. The oscillatory component was calculated by subtracting the aperiodic component from the total power and averaging over delta (1–4 Hz), theta (4–8 Hz), and alpha (8–11 Hz) bands.

### Blood sampling

Polyphasic participants underwent blood sampling during the 24-h in-depth monitoring lab visit before and towards the end of the polyphasic sleep schedule. Blood samples were collected via an indwelling venous catheter every 30 min for 24-h (from 16:00 on the first day till 16:00 on the next day). Venous access was established one hour before starting blood sampling to avoid cortisol peaks in the first two blood samples. Each blood sample was mixed up with 150 μl Trasylol and 150 μl EDTA and centrifuged. The extracted blood serum was collected in two 1-ml serum collection tubes and stored in the freezer until the analysis. Cortisol and melatonin concentrations were assessed by radioimmunoassay; free human growth hormone concentration was evaluated with the IMMULITE 2000 Systems Analyzers *in vitro* by an expert blind to the participant’s characteristics and sleep data.

### Cognitive tasks

During their two lab visits, polyphasic participants and controls performed cognitive tasks, including declarative memory (LGT-3, Bäumler, 1974), procedural memory (Shimoyama et al., 1990), fluid reasoning (BOMAT, Hossiep et al., 2001), and psychomotor vigilance (PVT, Dinges & Powell, 1985). Specifically, the fluid reasoning test assessed intellectual performance using matrix pictures with missing fields. The psychomotor vigilance test counted the number of lapses in attention. The declarative learning test evaluated training and overnight effects of visual recognition (city map, objects, and paired associations) and verbal memory (vocabulary, telephone numbers, figures, names, and abstract terms). The procedural memory test evaluated the training and overnight effects of the sequential finger-tapping task, with the training effect being calculated as the ratio between the first and the last evening scores, and the overnight consolidation effect being calculated as the ratio between the morning and first evening scores.

For the polyphasic participant, all tests were performed before and 5 weeks after the switch to the polyphasic sleep schedule. For the controls who stayed on the monophasic sleep schedule, cognitive performance was tested twice with a lag of 5 weeks between the test sessions. Changes in cognitive performance from the baseline period were calculated by dividing the scores from the second session (week 9) by the scores from the first session (week 4). The value of 1 of these ratio scores indicates no change in cognitive performance, whereas values higher and lower than 1 indicate that the second session showed higher and lower scores, that is better and worse performance, respectively.

### Psychological assessment

The polyphasic participant completed psychological questionnaires for 4 weeks before and 5 weeks after the switch to the polyphasic schedule once per week. The monophasic controls completed weekly psychological questionnaires for 9 weeks of the monophasic schedule. The presence and severity of manic/hypomanic and depressive symptoms were assessed with the Altman Self-Rating Mania Scale (ASRM, Altman et al., 1997) and the simplified Beck Depression Inventory (BDI-V, Schmitt et al., 2006), respectively. Higher scores reflected more manic/depressive symptoms. Subjective cognitive performance was assessed with the FLei questionnaire, which covers attention, memory, executive function, and visual neglect domains (Beblo et al., 2010). A higher score reflected worse subjective performance.

All questionnaire scores were baseline normalized by the average of the baseline period. Specifically, each one of the 9 scores corresponding to the 9 weeks of the experiment was divided by the mean of the scores of the first 4 weeks of the experiment (before the switch to the polyphasic sleep schedule for the participant) such that the ratios higher than one indicate higher scores after the switch.

### Statistical analysis

For different domains, we used different statistical approaches based on the data availability. For the endocrine data, we used the within-subject evaluation based on the visual inspection with no statistical test applied. This approach was forced by the fact that all but one participant terminated the polyphasic sleep schedule before the end of the experiment. Specifically, we compared the polyphasic and monophasic growth hormone, cortisol, and melatonin levels of the single participant who did adhere to the polyphasic sleep schedule.

For the EEG data, we compared spectral power-related variables using the two-tailed *t*-test as here, we had a relatively large number of 30-s epochs (see Table S1). Alpha-level was set at 0.05, the effect sizes were evaluated with Cohen’s d. Only the polyphasic variables that differed from both the corresponding monophasic variables of the same participant and controls were considered statistically significant.

Cognitive and neuropsychological characteristics of a single participant were compared to those of ten controls who stayed on the monophasic sleep schedule during the entire experiment using a modified *t*-test adapted from Crawford & Howell (1998). This method treats the individual as a sample of N = 1, being an appropriate approach to use when the normative sample is small.

The sleep architecture of the polyphasic participant was compared to that from the reference dataset (n = 97) using the modified *t*-test (Crawford & Howell, 1998).

For EEG, sleep stages, cognitive tests, and neuropsychological questionnaires, Benjamini-Hochberg’s adjustment was applied to control for multiple comparisons with a false discovery rate set at 0.05.

## Results

Nine out of ten participants terminated the polyphasic sleep schedule during the first three weeks of the experiment. The main reasons for the drop-out included a too strong impact on their professional or private life, adaptation length, daily naps, and cognitive burden (Fig. S1). One participant (a 25-year-old male) succeeded to adhere to the polyphasic schedule for 5 weeks with a median cumulative sleep time of only 3 h/day (range 2–10). The participant succeeded in strictly adhering to the *Uberman* schedule for 22/35 days of the experiment. For the other 13/35 days, he showed a schedule with an extended core sleep period of ≥4 hours, reportedly due to unintentional oversleeping. For comparison, the median total sleep time during the monophasic schedule was 7 h/day (range 4–8, Fig. 1A–B, Fig. S2).

Below we report the sleep, endocrine, cognitive, and affective characteristics of this participant before and after switching to the polyphasic sleep schedule.

Polyphasic and monophasic schedules presented comparable sleep architecture with only a slight (7%) decrease in relative REM sleep amount in favor of slow-wave sleep (SWS) in the polyphasic compared to the monophasic schedule (Fig. 1C–D). Also, as in monophasic sleep, REM sleep appeared more frequent after sleep episodes with higher SWS sleep surpassed earlier in the 24-hour cycle in the polyphasic schedule, showing that the general “REM after SWS” pattern has been conserved in some way. Neither the monophasic nor the polyphasic distribution of sleep stages of the participant was statistically different from that measured in the reference dataset (Fig. 1E). Nevertheless, probabilities of scoring certainty predicted by the machine learning algorithm were lower for the polyphasic compared to the monophasic sleep schedule of the same participant for the N2 and SWS stages (Fig.S3), suggesting less distinguishable sleep stages during polyphasic sleep.

Polyphasic REM sleep showed decreased delta, theta, and alpha-band oscillations and increased (i.e., flatter) low-band aperiodic slopes compared to both the own monophasic schedule and controls with medium to large effect sizes (Fig. 2, Table S1). During N2 and SWS, polyphasic spectral power was within the reference dataset range.

**Figure 2.**
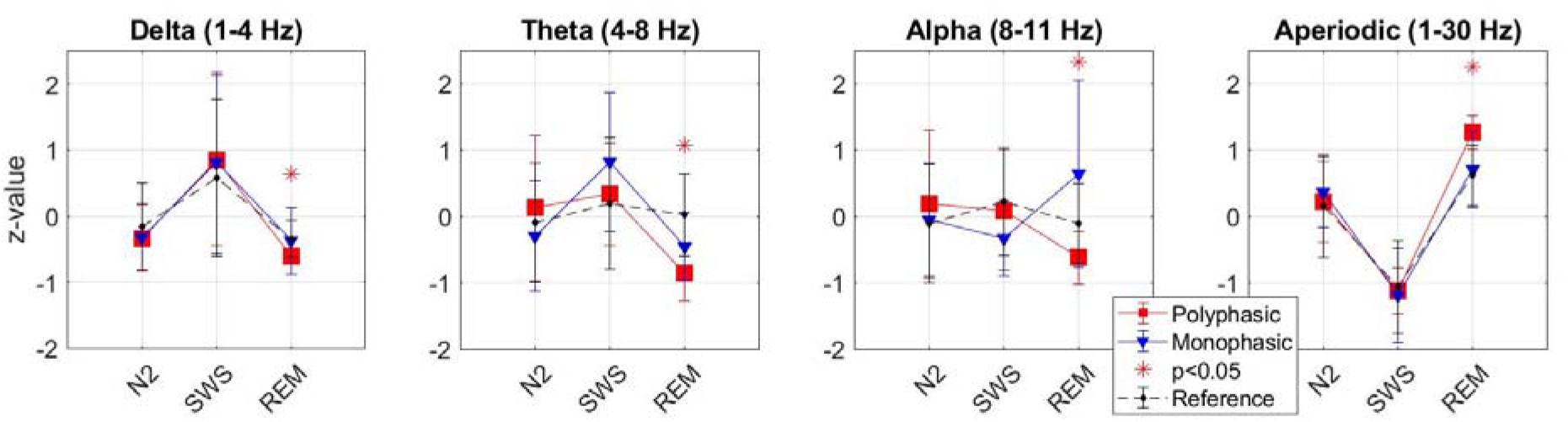
Spectral power during the monophasic and polyphasic sleep schedule. EEG power was averaged over F4, C4, and O2 electrodes and differentiated into its oscillatory and aperiodic components, z-scores transformed, and averaged over each sleep stage of the monophasic (blue lines with triangles) and polyphasic (red lines with squares) sleep schedules of the same participant. In addition, the same variables are shown for the age-matched controls (black dashed lines with diamonds) from the reference dataset (n = 97). Error bars denote standard deviations between the 30-s epochs. The oscillatory power component in the delta, theta, and alpha frequency bands as well as the slopes of the aperiodic power component in the low (1–30 Hz) frequency band are shown separately. Polyphasic REM sleep shows decreased delta, theta, and alpha-band power as well as flatter (more positive) aperiodic slopes compared to own monophasic sleep and controls (marked with the asterisks, see also Table S1). All spectral power characteristics during N2 and SWS are comparable. REM: rapid eye movement, N2: non-rapid eye movement stage 2, SWS: slow-wave sleep.

After 5 weeks of polyphasic sleep, the overall release of growth hormone appeared strongly decreased. Specifically, the area under the curve (AUC) decreased by 95% (from 1300 to 67 ng/ml per day). Moreover, the pattern of its secretion had changed: instead of one major peak during the first hours of sleep (8 ng/ml), six minimal peaks (0.2 ng/ml) were observed after each sleep episode, reflecting a decrease of 97.5%. Cortisol and melatonin secretion was comparable between the two test sessions (Fig. 3).

**Figure 3.**
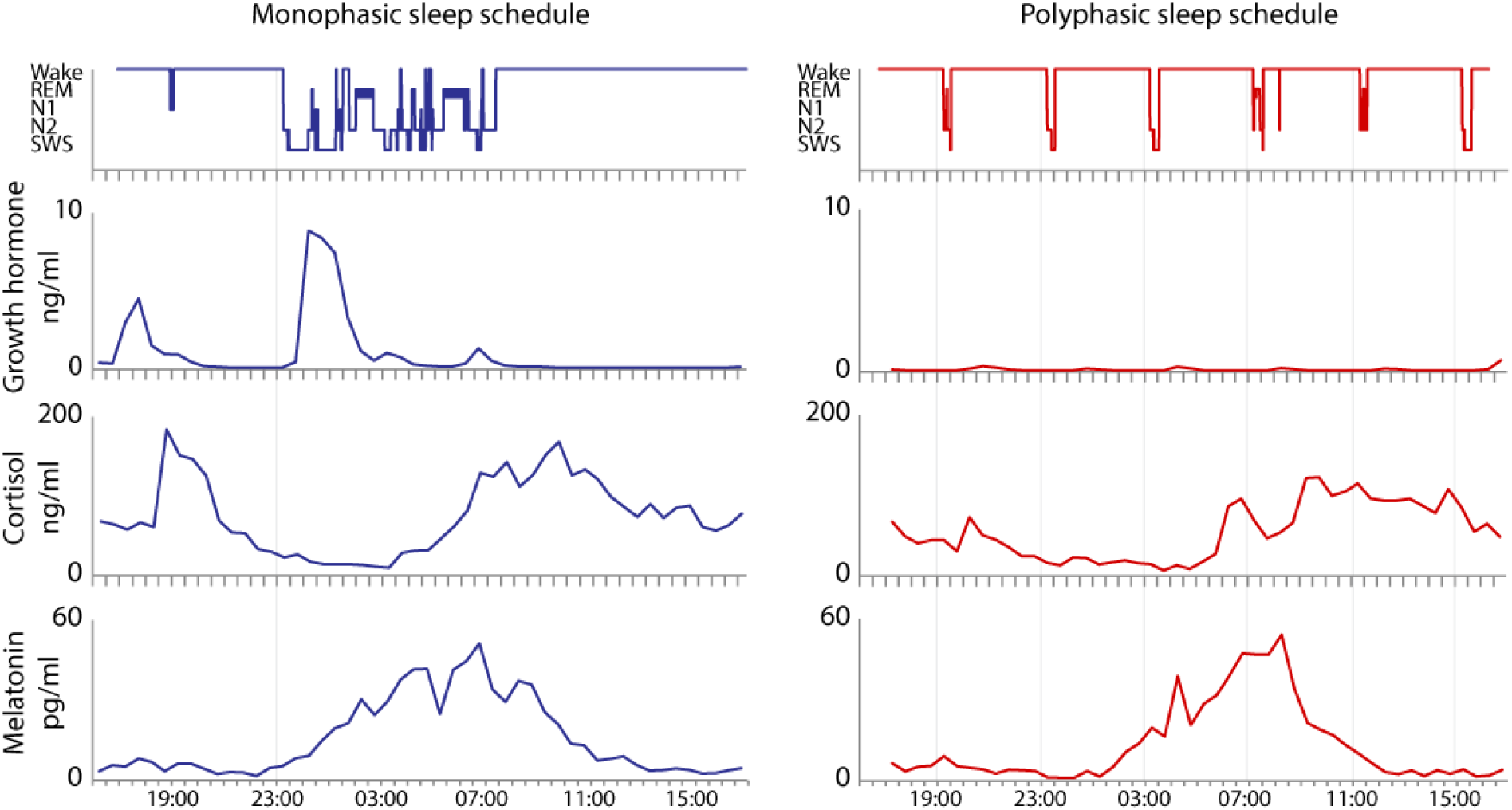
Monophasic and polyphasic endocrine profile. From the upper panel to the lower, growth hormone, cortisol, and melatonin levels, respectively, are depicted over 24 hours, time-locked to the hypnograms (top panel) of the monophasic (left) and polyphasic sleep (right) within the same participant. Light gray lines indicate sleep onset. While cortisol and melatonin secretion was comparable between the two test sessions, growth hormone release was strongly suppressed during the polyphasic compared to the monophasic sleep schedule. Six small peaks can be observed after each nap (right) instead of one major peak during the first hours of sleep (left).

After 5 weeks of polyphasic sleep, the participant did not show major changes in cognitive performance, depressive symptoms or subjective performance compared to controls (Fig. 4). In the psychomotor vigilance task, the participant showed a 1.75-fold increase in the number of lapses in attention compared to baseline; however, after controlling for multiple comparisons, this increase was not statistically different from that measured in controls. During the first week of the polyphasic sleep schedule, the participant showed a significant increase in manic symptoms compared to controls (t = 2.89, p = 0.008 < corrected p-value = 0.01); however, this difference diminished from the second week till the end of the experiment (Fig. 4B).

**Figure 4.**
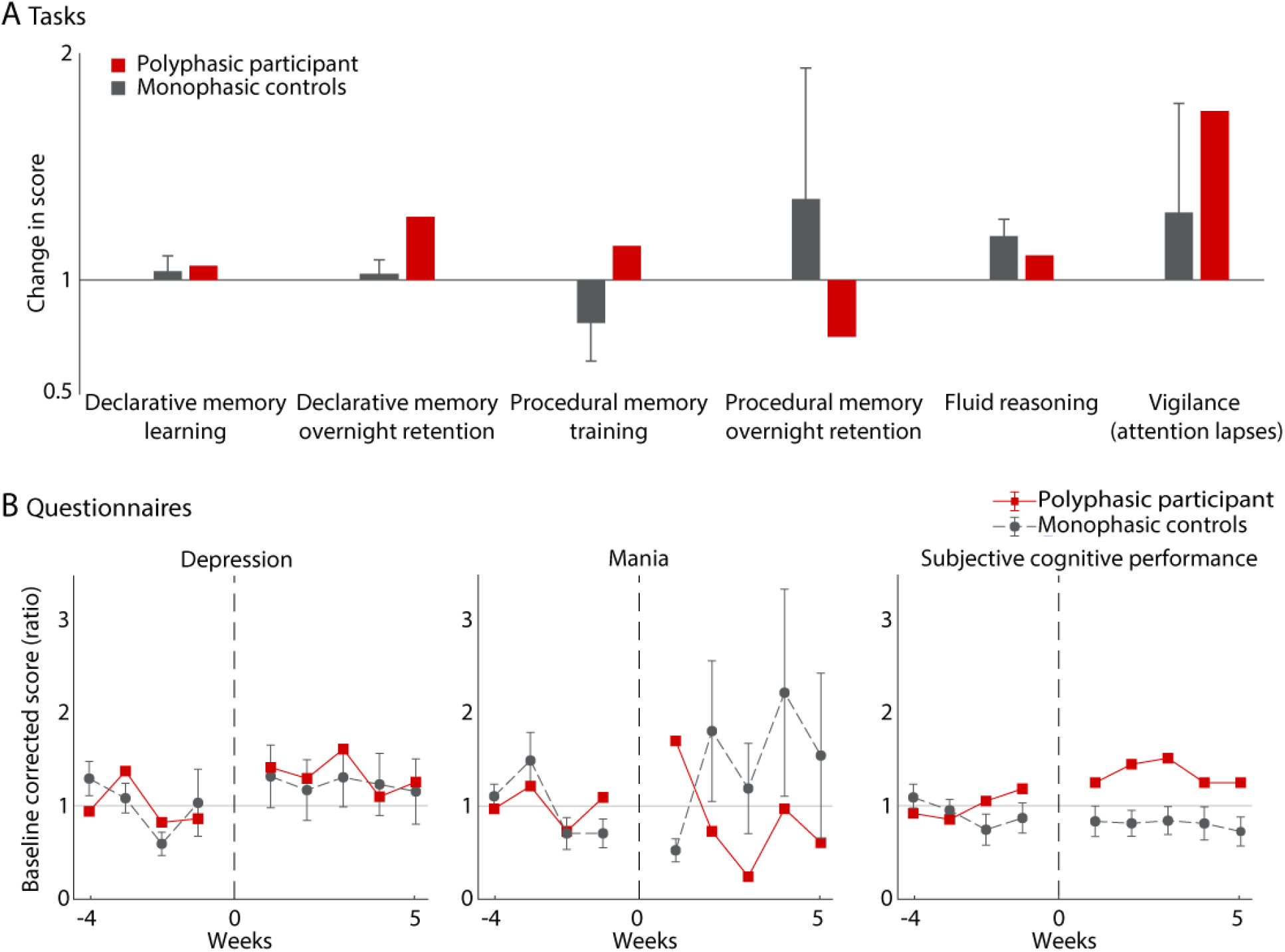
Cognitive and affective profile during the polyphasic and monophasic sleep schedules. (A) Cognitive performance in the domains of vigilance, declarative memory, fluid reasoning, and procedural memory are depicted as the ratios between the scores after and before the switch to the polyphasic sleep schedule for the individual adhering to the polyphasic schedule for the entire study period (red bars) and for 10 control participants who completed the same measurements but remained on a monophasic sleep schedule (gray bars). The ratios>1 indicate higher scores in the second session (after switching to the polyphasic schedule for the polyphasic participant). Error bars denote standard errors. After 5 weeks of polyphasic sleep, the participant did not show any changes in fluid reasoning and immediate or overnight effects of declarative and procedural memory as reflected by the ratios close to one. In the psychomotor vigilance task, the participant showed a 1.75-fold increase in the number of lapses in attention compared to baseline, however, this increase was not statistically different from that measured in controls. (B) Baseline-normalized scores for the questionnaires measuring depression, mania, and subjective cognitive performance are shown for 4 weeks before and 5 weeks after switching to the polyphasic schedule for the polyphasic participant (red line with squares), and for the same length for 10 control participants remaining on a monophasic sleep schedule (gray dashed line with rounded squares). The vertical dashed line marks the switch from the monophasic to the polyphasic sleep schedule for the participant. During the 5 weeks of the polyphasic sleep schedule (weeks 1 to 5), the participant did not show any changes in depression and subjective performance compared to the scores before the switch (weeks -4 to -1) as reflected by the ratios being less than 1.5. The participant showed a significant increase in manic symptoms compared to controls during the first week of the polyphasic schedule (marked with an asterisk), which disappeared afterward. Albeit not statistically different from controls, the polyphasic participant reported worse subjective cognitive performance (higher scores indicate worse performance) after switching to the polyphasic schedule.

## Discussion

We aimed to explore the effect of the sustained polyphasic sleep schedule on the neural, endocrine, and cognitive profile of healthy young students. The study was initiated by the participants, who out of intrinsic motivation approached the research team and invested considerable time and effort without payment, thus demonstrating a very high degree of commitment. On this background, our study had a striking drop-out rate of 90%, indicating that the impact of the polyphasic sleep schedule on participants’ lives was mostly intolerable despite strong intrinsic motivation to persist. Only one volunteer succeeded in adhering to the polyphasic schedule for five weeks and participating in all planned data acquisition.

We found that polyphasic sleep was associated with strongly decreased growth hormone – but not cortisol or melatonin – levels. Neurophysiological patterns of the polyphasic schedule included decreased delta, theta, and alpha-band oscillatory and increased (i.e., flatter) aperiodic activity during REM sleep. Sleep stage proportions, mood, and cognitive performance during the polyphasic sleep schedule were comparable to those measured during the monophasic schedule except for a tendency towards decreased psychomotor vigilance and a higher pressure for SWS over REM sleep observed during the polyphasic schedule. Below we discuss each finding in detail.

Regarding cognitive performance, we found that the participant tended to experience more lapses in attention during the polyphasic compared to the monophasic schedule. This can reflect reduced cortical responsiveness to incoming stimuli due to the observed alterations in sleep EEG activity. Quite surprisingly, the participant showed no other cognitive alterations, namely, no decline in fluid reasoning, declarative, and procedural memory. This suggests that a polyphasic schedule does not interfere cardinally with the cognitive function of sleep, at least at the single-case level. A possible explanation is that the brain has a mechanism for adaptation to chronic sleep restriction to stabilize performance (Belenky et al., 2003) and thus provide evolutionary survival benefits during demanding conditions that involve sleep reduction (e.g., lactation, natural disasters). Such a mechanism might, however, have somewhat limited capacity, operating successfully only on the short-term and/or middle-term but not long-term scale.

Similarly, the affective characteristics of the participant did not change drastically during the five weeks of the polyphasic schedule. His scores on the depression self-questionnaire remained on the same level as before the switch to the polyphasic schedule and his scores on the mania self-questionnaire showed only a transient increase during the first week of the polyphasic schedule, which disappeared afterward. Given that sleep is crucial for the regulation of emotional processes, mood, and affective well-being (Dresler et al., 2014), our observation is rather unexpected. Notably, disrupted sleep reported in some patients with major depressive disorder has been related to the disintegration of the internal connections between sleep, affective, cognitive, and other physiological cycles, such as melatonin and cortisol concentrations (Hickie and Rogers, 2011).

Overall, our findings contrast with the literature reporting that polyphasic sleep is associated with mood deterioration, emotional discomfort, irritability, and increased depression ratings as a function of the duration of exposure to polyphasic sleep (Weaver et al., 2021). It should be kept in mind, however, that the level of self-assessed emotional impairments of polyphasic sleepers can be under or overestimated due to sleep deprivation (Weaver et al., 2021).

The most prominent feature of polyphasic sleep observed in our study was a strong overall suppression of growth hormone (GH). Moreover, we found that the pattern of GH secretion had changed: instead of one major peak during the first SWS episode, we observed six minimal peaks after each nap. This is in line with the study in young men reporting that the normal nocturnal GH surge disappears with sleep deprivation (Davidson et al., 1991). For regular sleep, the GH surge coincides with the first stage of SWS (Steiger, 2007; Dresler et al., 2014). Moreover, both GH and SWS are likely promoted by the same substance, the growth hormone-releasing hormone (reviewed in Steiger, 2007; Dresler et al., 2014). After sleep deprivation, GH secretion during sleep is prolonged and an additional GH peak (not associated with SWS) is observed in some but not all volunteers, indicating less reliable temporal associations between GH and SWS (Davidson et al., 1991).

Importantly, besides stimulating the growth of bones and muscles and intermediate metabolism, GH also has some neuroprotective effects, such as a decrease in neuronal apoptosis due to oxidative stress (Åberg et al., 2006). Moreover, GH improves mood, motivation, working capacity, and cognition and modulates learning and memory, possibly through its effects on the monoaminergic, dopaminergic, glutamatergic, and cholinergic systems (Åberg et al., 2006). In line with this, it has been reported that adults with GH deficiency who do not receive replacement therapy usually show significant psychological alterations, such as lack of energy, impaired memory, and other cognitive alterations (Wass & Reddy, 2010). Correspondingly, GH administration improves self-reported well-being as well as short and long-term memory in GH-deficient young adults and probably in elderly people (Åberg et al., 2006).

This literature is at odds with our findings on the preserved cognitive performance during the polyphasic sleep schedule. One should keep in mind, however, that GH effects reported in the literature occur on a much longer time scale (months, years) than those reported here (five weeks). Striving to adopt a habitual polyphasic schedule on a longer scale (months, years) might eventually adversely affect cognitive performance and affective well-being. In addition, we should emphasize that we explored the polyphasic GH release at the ligand level only. It is possible that at the receptor level, some regulatory changes compensated for the decreased GH release. Interestingly, polyphasic sleep has not shown any changes in cortisol or melatonin levels, suggesting a strong circadian regulation of these hormones.

Neurophysiological activity after five weeks of polyphasic sleep was characterized by decreased delta, theta, and alpha-band oscillations during REM sleep. In the literature, decreased alpha oscillations were observed after the prolonged wake due to acute sleep deprivation (Wu et al., 2021), suggesting that polyphasic sleep restriction builds up chronic sleep deprivation. Moreover, we found that polyphasic REM sleep showed flatter slopes of the aperiodic power component (i.e., slower decay of the power spectrum) compared to monophasic REM sleep. It has been suggested that flatter high-band aperiodic slopes might reflect a shift in the excitation-to-inhibition ratio in favor of excitation (Gao et al., 2017; Lendner et al., 2020; Miskovic et al., 2019). Moreover, recent findings have revealed that flatter aperiodic activity is associated with higher peripheral levels of amyloid-beta, a toxic metabolite accumulating during wakefulness (Rosenblum et al., 2023 b).

Interestingly, polyphasic sleep showed rather preserved sleep architecture with a classical distribution of sleep stages: the sleep episodes of the early night hours presented with SWS, whereas the sleep episodes of the morning presented with REM sleep (Fig. 1D). This points to the high level of plasticity of sleep and circadian regulatory mechanisms. A possible interpretation is that when on a polyphasic sleep schedule, the brain relies more on ultradian oscillators, which normally cycle in concert with the circadian clock. An example of such an oscillator reported in mice would be the dopaminergic ultradian oscillator that generates bursts of activity and alertness every several hours, regulating arousal (Blum et al., 2014). Nevertheless, the certainty of the sleep stages predicted by the machine learning algorithm was lower for the polyphasic compared to the monophasic sleep schedule (Fig.S2). This means that sleep stages are less distinguishable in polyphasic sleep, possibly indicating that sleep stages are condensed due to high sleep pressure and decreased sleep duration.

The sleep deficiency that is inherent to polyphasic sleep schedules is a major concern as it is known to affect multiple aspects of physical and mental health, performance, and safety (McCoy et al., 2011; Weaver et al., 2021). Even though we observed preserved sleep architecture, an unavoidable decrease in the absolute amount of SWS raises a serious concern regarding e.g. glymphatic brain clearance: it has been shown that even one night of sleep deprivation affects the amyloid-beta burden in the hippocampus of healthy volunteers (Shokri-Kojori et al., 2018). A prolonged reduction of the SWS amount may, therefore, severely interfere with brain clearance, leading to the accumulation of toxic metabolites, such as amyloid-beta, thereby increasing the risk of eventual cognitive impairment and neurodegeneration (Ju et al., 2014). Nevertheless, to date, no evaluations of the long-term (months, years) impact of polyphasic sleep restriction (and an unavoidable reduction of the overall amount of SWS bound to this schedule) have been done (Weaver et al., 2021) and probably will not be done due to significant ethical concerns in terms of neurobiological costs, which can accumulate over time.

The major limitation of this study is that it compares monophasic and polyphasic sleep within a single participant and, therefore, does not allow us to generalize the observed effects to the broader population. Nevertheless, the fact that our participant showed comparable sleep architecture and spectral power to that recorded in a large group of age-matched controls suggests that observed findings are only based on physiological idiosyncrasies of the participant. Moreover, the pre-study screening has shown that the participant was healthy with no family history of any sleep disorders. He had no night shifts before or during the experiment and experienced the polyphasic schedule for the first time. Therefore, this participant could be seen as representing a typical individual from a young population.

A fact to keep in mind, however, is the existence of considerable interindividual variability in the homeostatic response to sleep deprivation (Rusterholtz et al., 2016). Variability in this response can partially explain why only one out of ten participants of this study could adhere to the polyphasic schedule preserving relatively unimpaired cognitive and affective characteristics whereas the rest of the group could not remain on such a schedule for more than several days (i.e., survivorship bias). Accordingly, polyphasic sleep schedules might be more detrimental to some people than others.

Likewise, during the polyphasic schedule, the participant had two days with unintentional recovery sleep periods of 10 hours as he did not hear the alarm clock (Fig. S2). During the test day, he strictly adhered to the polyphasic rhythm, accumulating a total of 2 hours of sleep.

Despite these limitations, this study is highly relevant from both basic scientific and clinical points of view. It clearly shows that on one hand, a healthy young man can survive on a polyphasic schedule for at least five weeks with a mostly preserved sleep architecture and very few cognitive and affective changes. On the other hand, radically polyphasic sleep severely affects growth hormone release, and, therefore, interferes with at least one (endocrine) function of sleep. Considering the neurophysiological changes, decreased vigilance, and abolished growth hormone release associated with the polyphasic schedule as well as decreased absolute time spent in REM and SWS, it is doubtful that radically polyphasic sleep schedules can subserve the different long-term functions of sleep to a sufficient degree.

## Contributors

MD designed the study. MR, NK, BB, MZ, MD acquired the data. ZZ, YR, FDW, MR, BB, MU, MD analyzed the data. YR wrote the manuscript. ZZ, YR, MD designed the figures. AS, MD acquired the funding. All authors discussed and interpreted the data and contributed to, reviewed, revised, and approved the final draft of the paper. All authors had full access to all the data in the study and had final responsibility for the decision to submit for publication.

## Acknowledgments

We would like to thank the involved polyphasic sleepers for their dedication to the preparation and execution of the study. This work was supported by the Max Planck Institute of Psychiatry, Munich, and a Vidi fellowship by the Dutch Research Council (NWO).

## Data sharing

Deidentified data is available to researchers upon request from the corresponding authors.

## Abbreviations

GH: growth hormone
REM: rapid eye movement
SWS: slow-wave sleep

