## Supplementary Material for "Sustained polyphasic sleep restriction abolishes human growth hormone release"

Yevgenia Rosenblum* [1], Frederik D. Weber* [1,2], Michael Rak* [3], Zsófia Zavecz* [4],

Nicolas Kunath [3], Barbara Breitenstein [3], Björn Rasch [5], Marcel Zeising [6], Manfred Uhr [3],

Axel Steiger [3], Martin Dresler [1]

[1] Radboud University Medical Centre, Donders Institute for Brain, Cognition and Behavior, Nijmegen, Netherlands, [2] Netherlands Institute for Neuroscience, Department of Sleep and Cognition, Amsterdam, Netherlands, [3] Max Planck Institute of Psychiatry, Munich, Germany, [4] Center for Human Sleep Science, Department of Psychology, University of California Berkeley, Berkeley, California, [5] Department of Psychology, Division of Biopsychology, University of Zurich, Zurich, Switzerland, [6] Klinikum Ingolstadt, Centre of Mental Health, Ingolstadt, Germany

* - equal contribution

**Corresponding authors:**

Yevgenia Rosenblum, Radboud University Medical Centre, Donders Institute for Brain, Cognition and Behavior, Kapittelweg 29, 6525 EN Nijmegen, the Netherlands.

Martin Dresler, Radboud University Medical Centre, Donders Institute for Brain, Cognition and Behavior, Kapittelweg 29, 6525 EN Nijmegen, the Netherlands.

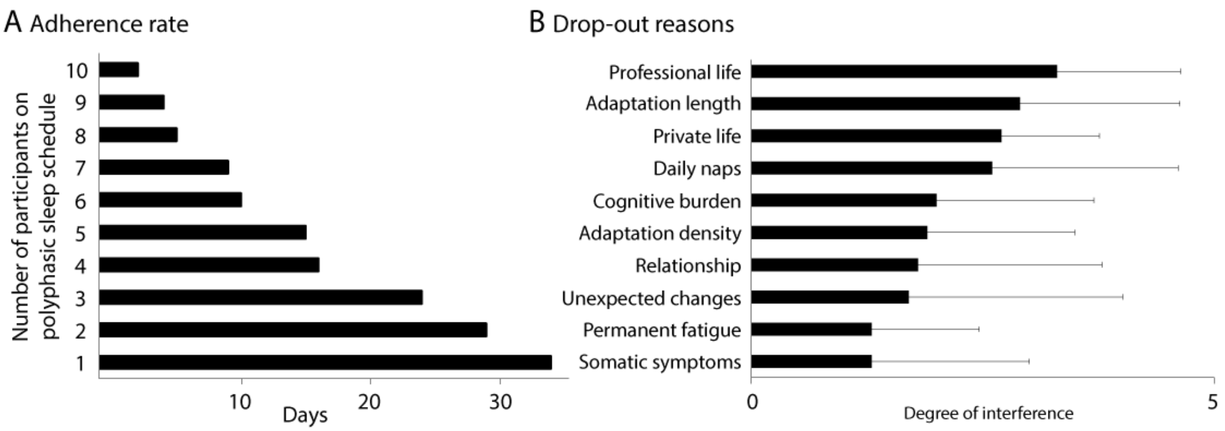


**Figure S1. Drop-out of the study.** All but one of the test subjects terminated the polyphasic sleep schedule during the first three weeks (A), mainly due to a too strong impact on their social lives. The reasons which led to the dropout of the polyphasic sleep schedule were assessed with the self-questionnaire, which evaluated the degree of interference of each parameter on a scale from 1 to 6 (B). Error bars denote standard deviations.

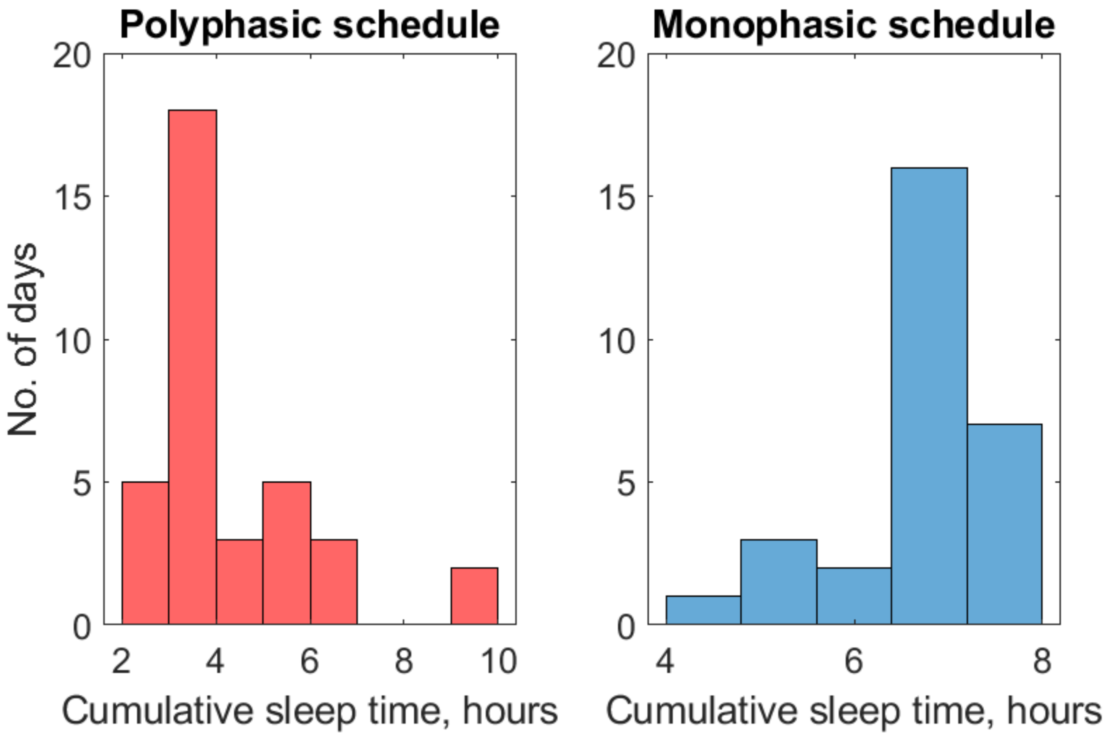


**Figure S2. Cumulative sleep time during monophasic and polyphasic sleep schedules.** Histograms of the cumulative sleep time before (right) and after (left) switching to the polyphasic sleep schedule. During the polyphasic period, the participant succeeded in strictly adhering to the *Uberman* schedule for 22/35 days of the experiment.

**Table S1:** Statistical comparison of monophasic and polyphasic power

| Stage | No. of epochs | | Oscillatory component | | | | | | Aperiodic component | | | |
|---|---|---|---|---|---|---|---|---|---|---|---|---|
| | | | Delta | | Theta | | Alpha | | Low band | | High band | |
| | poly | mono | p | d | p | d | p | d | p | d | p | d |
| N2 | 111 | 412 | 0.768 | 0.032 | <0.001 | -0.476 | 0.013 | -0.266 | 0.014 | 0.262 | 0.015 | 0.260 |
| SWS | 72 | 250 | 0.843 | -0.027 | <0.001 | 0.494 | <0.001 | -0.645 | 0.417 | -0.109 | 0.001 | -0.435 |
| REM | 35 | 167 | **0.006** | 0.514 | **<0.001** | 0.813 | **<0.001** | 0.978 | **<0.001** | -1.068 | 0.545 | -0.113 |

Monophasic and polyphasic sleep EEG patterns were compared using the t-test, effect sizes were assessed with the Cohen's d. Bold font marks the polyphasic variables that were statistically significantly different compared to both the own monophasic sleep as well as the aged-matched controls from the reference dataset (Figure 2). N: non-REM, SWS: slow-wave sleep, REM: rapid eye movement, epoch: 30 seconds.

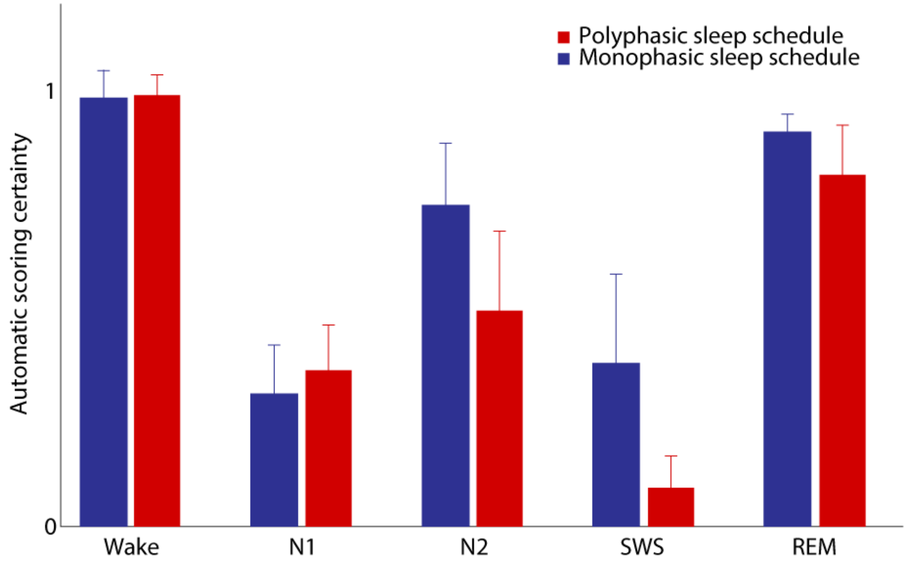


**Figure S3. Scoring certainty of the monophasic and polyphasic sleep schedules.** Scoring certainty for each sleep stage by an automatic algorithm (Abou Jaoude et al., 2020) for the polyphasic (red bars) and monophasic (blue bars) schedules. Error bars denote standard deviations. The probabilities predicted by the machine learning algorithm are lower for the polyphasic compared to the monophasic sleep schedule for the N2, SWS and REM stages, possibly indicating that sleep stages are condensed due to the high sleep pressure and decreased sleep durations. REM: rapid eye movement, N2: non-rapid eye movement stage 2, SWS: slow-wave sleep.
